# Timing Determines Tuning: a Rapid Spatiotemporal Transformation in Superior Colliculus Neurons During Reactive Gaze Shifts

**DOI:** 10.1101/302125

**Authors:** Morteza Sadeh, Amirsaman Sajad, Hongying Wang, Xiaogang Yan, John Douglas Crawford

**Affiliations:** York Centre for Vision Research and Vision: Science to Applications Program; York Neuroscience Graduate Diploma Program; Canadian Action and Perception Network (CAPnet); Departments of Psychology, Biology, and Kinesiology and Health Science, York University, Toronto, Ontario, Canada M3j 1P3

## Abstract

Gaze saccades –rapid shifts of the eyes and head toward a goal— have provided fundamental insights into the neural control of movement. For example, it has been shown that the superior colliculus (SC) transforms a visual target (T) code to future gaze (G) location commands after a memory delay. However, this transformation has not been observed in ‘reactive’ saccades made directly to a stimulus, so its contribution to normal gaze behavior is unclear. Here, we tested this using a quantitative measure of the spatial continuum between T and G coding based on variable gaze errors. We demonstrate that a rapid T-G transformation occurs between SC visual and motor responses during reactive saccades, even *within* visuomotor cells, with a continuous spatiotemporal shift in coding occurring in cell types (visual, visuomotor, motor). We further show that the primary determinant of this spatial code was not the intrinsic visual-motor index of different cells or populations, but rather the *timing* of the response in *all* cells. These results suggest that the SC provides a rapid spatiotemporal transformation for normal gaze saccades, that its motor responses contribute to variable gaze errors, and that those errors arise from a noisy spatiotemporal transformation involving all SC neurons.

**Significance Statement:** Oculomotor studies have demonstrated visuomotor transformations in structures like the superior colliculus with the use of trained behavioral manipulations, like the memory delay and antisaccades tasks, but it is not known how this happens during normal saccades. Here, using a spatiotemporal model fitting method based on endogenous gaze errors in ‘reactive’ gaze saccades, we show that the superior colliculus provides a rapid spatiotemporal transformation from target to gaze coding that involves visual, visuomotor, and motor neurons. This technique demonstrates that SC spatial codes are not fixed, and may provide a quantitative biomarker for assessing the health of sensorimotor transformations.

## Introduction

Saccades and rapid gaze shifts involving coordinated eye-head motion have been employed extensively to study the fundamental neural basis of sensorimotor transformations (Mays and Sparks 1980, Wurtz and Albano 1980, Gnadt, Bracewell et al. 1991, Deubel 1995, Freedman and Sparks 1997, Freedman and Sparks 1997, Freedman 2008, Sadeh et al. 2015, Sajad et al. 2015). As a result, the circuitry of the saccades system in humans is very well described (Fischer 1986, Pierrot-Deseilligny et al. 1991, Pierrot-Deseilligny, Rivaud et al. 1991, Gaymard and Pierrot-Deseilligny 1999, Munoz and Everling 2004). Studies in non-human primates have revealed numerous additional details about the cellular and signal properties. For example, neurons with gaze-related responses in the superior colliculus (SC), frontal eye fields, (FEF) and lateral intraparietal cortex (LIP) can be categorized into populations of cells with ‘visual’ responses (briefly delayed burst responses to a visual stimulus), ‘motor’ responses (burst activity just before and after a saccade) or visuomotor responses, i.e., both visual and motor (Goldberg and Wurtz 1972, Goldberg and Wurtz 1972, Harris 1980, Goldberg and Bushnell 1981, Bruce and Goldberg 1985, Bruce, Goldberg et al. 1985, Munoz and Wurtz 1995, Munoz and Wurtz 1995, Freedman and Sparks 1997, Bisley and Goldberg 2003, Gandhi and Katnani 2011). The timing of these responses seems to imply a spatiotemporal transformation between the visual and motor responses, but demonstrating this transformation in the spatial domain is not trivial.

Normally there is little or no temporal separation between visual and motor responses, and little separation between the direction of a visual stimulus and saccade direction, so visual and motor responses are easily conflated in both the temporal and spatial domains. The technical challenge for spatial separation is that the key parameters –retinal location of a visual target and gaze displacement— only diverge in the presence of saccade errors (Mays and Sparks 1980, Waitzman, Ma et al. 1988, Stanford and Sparks 1994, Munoz and Everling 2004), ocular torsion (Crawford and Guitton 1997, Klier and Crawford 2003), or very large gaze shifts (Klier, Henriques et al. 2002). Studies of this question have mainly used saccade errors and focused on structures such as the midbrain superior colliculus (SC) and cortical areas like the frontal eye fields (FEF), and lateral intraparietal cortex (LIP). In general, many experiments suggest that these structures employ a retinal spatial code (Klier, Wang et al. 2001, Martinez-Trujillo, Medendorp et al. 2004, Avillac, Deneve et al. 2005, Constantin, Wang et al. 2007, DeSouza, Keith et al. 2011, Monteon, Wang et al. 2013, Sadeh, Sajad et al. 2015, Sajad, Sadeh et al. 2015), although some have suggested they encode displacement of gaze direction (Mays and Sparks 1980, Freedman and Sparks 1997, Horwitz and Newsome 1999, Knight and Fuchs 2007, Marino, Rodgers et al. 2008). A use of a purely retinal code would seem to suggest that the conversion into motor coordinates only happens further downstream, in structures such as the brainstem reticular formation (Sparks 1989, Sparks and Hartwich-Young 1989, Snyder 2000, Sparks 2002, Crawford, Henriques et al. 2011, Sadeh, Sajad et al. 2015).

The challenge for detecting a spatiotemporal transformation is even higher, because it requires distinguishing retinal and motor codes within the short time span of the neural response to a single saccade target. One useful technique is to train animals to saccade opposite to the target (the anti-saccade task), using a spatial dissociation between target position and gaze direction. This has shown that many cells in the SC, FEF, and LIP initially encode visual target direction, but then switch to coding saccade direction (Gnadt, Bracewell et al. 1991, Groh and Sparks 1992, Optican 1995, Gottlieb and Goldberg 1999, Russo and Bruce 2000, Marino, Rodgers et al. 2008,). Another approach is to separate visual and motor responses in time, through the interposition of a memory delay, and then fit various models against the response to targets at various directions in the presence of small, variable saccade errors. This showed that the SC and FEF visual response encodes target location (T) relative to the eye, whereas the motor response encodes future gaze direction (G) relative to the eye (Sadeh, Sajad et al. 2015, Sajad, Sadeh et al. 2015). A further spatiotemporal analysis of these results showed that the T-G transformation occurred continuously through intermediate codes during the memory delay, and then shifted to G in purely motor cells active just before a saccade (Sajad, Sadeh et al. 2016).

These findings, while important, employ experimental manipulations of behavior that are not normally present in saccades. For example, we do not normally look away from stimuli; this requires suppression signals and might cause the brain to imagine a target in the opposite direction (Bell, Everling et al. 2000, Everling and Munoz 2000, Munoz and Everling 2004, Coe and Munoz 2017). Likewise, we do not always delay saccades, and this task introduces suppression signals, memory signals, and a memory-motor transformation. These might introduce the accumulation of internal errors that artificially create an apparent ‘transformation’ (Gnadt, Bracewell et al. 1991, Stanford and Sparks 1994, White, Sparks et al. 1994, Ohbayashi, Ohki et al. 2003, Barber, Caffo et al. 2013, Hollingworth 2015, Sadeh, Sajad et al. 2015, Sajad, Sadeh et al. 2015, Sajad, Sadeh et al. 2016). Thus, it is not trivial to transpose these results to simply ‘reactive’ saccades made immediate and directly to a transient stimulus. It is simply not known whether a spatiotemporal transformation occurs during reactive saccades, and if so, how different SC cell types contribute to this transformation.

In the current study we directly investigated if the continuous neural activity present during reactive saccades shows the same spatial transformation that has been shown in the memory delay paradigm. To do this, we recorded from the same SC neurons using both the reactive and memory delay tasks, and analyzed their spatial content using a model fitting approach that we have developed and used recently (Keith and Crawford 2008, DeSouza, Keith et al. 2011, Sadeh, Sajad et al. 2015, Sajad, Sadeh et al. 2015). Further, we used a variant of our recent spatiotemporal analysis (Sajad, Sadeh et al. 2016) to test for a rapid transformation within the continuous burst present during reactive saccades. We found that, in the absence of a memory delay, SC neurons produce a rapid spatiotemporal transformation from retinal to gaze coding through a distributed transformation that appears to depend more on timing than cell type.

## Methods

### Animals and Surgical Procedures

The data were collected from two female monkeys (Macaca Mulatta, M1 and M2; age, 10 years; weights, 6.5 and 7 kg) with a protocol approved by the York University Animal Care Committee in accordance with guidelines published by the Canadian Council for Animal Care. With similar surgical procedures as described previously (Crawford, Ceylan et al. 1999, Klier, Wang et al. 2001), the monkeys were prepared for long-term electrophysiology and 3D gaze movement recordings. Each monkey was subjected to general anesthesia with 1–2% isoflurane after intramuscular injection of ketamine hydrochloride (10 mg/kg), atropine sulphate (0.05 mg/kg), and acepromazine (0.5 mg/kg). In order to minimize the collisions between experimental setup and Microdrive/electrode we implanted a vertically aligned unit recording chamber (i.e. with no tilt) placed 5 mm anterior and 0 mm lateral in stereotaxic coordinates, which allowed access to the left and right SC. This chamber angle and position were chosen to minimize collisions between the electrode/microdrive and the experimental setup during head movements, and to simplify the use of stereotaxic coordinates during recordings. The chamber was then surrounded by a dental acrylic cap, which was anchored to the skull with 13 stainless steel cortex screws. Two scleral search coils (diameter, 5 mm) were implanted in one eye of the monkeys to record 3D eye movements. Two orthogonal coils, which were secured with a screw on a plastic base on the cap, recorded the 3D head movements during the experiments. 3D recordings and analysis were performed as described previously (Crawford, Ceylan et al. 1999, DeSouza, Keith et al. 2011).

### Experimental equipment

We used a Pentium IV PC and custom-designed software to present stimuli, control behavior paradigms, send digital codes to a Plexon data acquisition system, and deliver juice rewards to the monkeys. Stimuli were presented on a screen 60 cm in front of the monkey, by use of a projector (WT600 DLP projector; NEC). Monkeys were seated on a custom-designed primate chair to have their heads move freely at the centre of a 1-m3 magnetic field generator (Crawford et al., 1999), and a juice spout (Crist Instruments) was placed on the skull cap for computer-controlled delivery of the juice reward to the monkey’s mouth.

### Behavioural recordings and paradigms

All experiments were performed in head-unrestrained conditions. This was necessary for the preliminary general reference frame analysis that preceded this experiment (Sadeh, Sajad et al. 2015). Here, target (T) and gaze (G) position in eye coordinates were the key parameters, but head unrestrained recordings also had advantages here: comfort, natural system behavior, adequate range of gaze motion for testing large neural response fields (RF; see below), and the tendency toward more prolonged neural activity for a spatiotemporal analysis (Keith, DeSouza et al. 2009, DeSouza, Keith et al. 2011). Conversely, 3D recordings and analysis were required for the proper transformation of T and G data to eye coordinates, to account for the significant torsional eye rotation and prominent non-linearities that occur in the head unrestrained gaze range (Tweed and Vilis 1987, Crawford, Ceylan et al. 1999, Klier, Wang et al. 2003, DeSouza, Keith et al. 2011).

The primary behavioral condition used during our neural recordings was the *Reactive* gaze shift task (Figure 1). The spatial aspects of this task were optimized for the model fitting analysis described below, including the separation of different reference frames and more importantly here, T from G coding. Animals were trained to begin each trial by fixating a central position (green circle with radius of 0.5°), with a location that randomly varied within a predetermined square range approximately equal to the cell’s RF size - for 900-1000 ms (randomly varied interval); simultaneous with initial fixation point disappearance-serving as GO signal - a target (red circle with a size of 0.5°) was presented in the periphery for 125 ms, brief enough to ensure no visual feedback after the completion of the gaze shift. The location was previously determined from preliminary RF mapping. Animals were then required to make a gaze shift toward the briefly flashing stimulus and fixate on it for 200 ms in order to receive juice reward. To spatially separate targets from gaze coding, we designated a tolerance window of 6–12° (diameter) for gaze errors around the locations of the targets, which resulted in a naturally-generated distribution of gaze end points around the targets (See Figure 1A, B, C). This variable error is the basis of our analysis method (Sajad, Sadeh et al. 2015).

**Figure 1:**
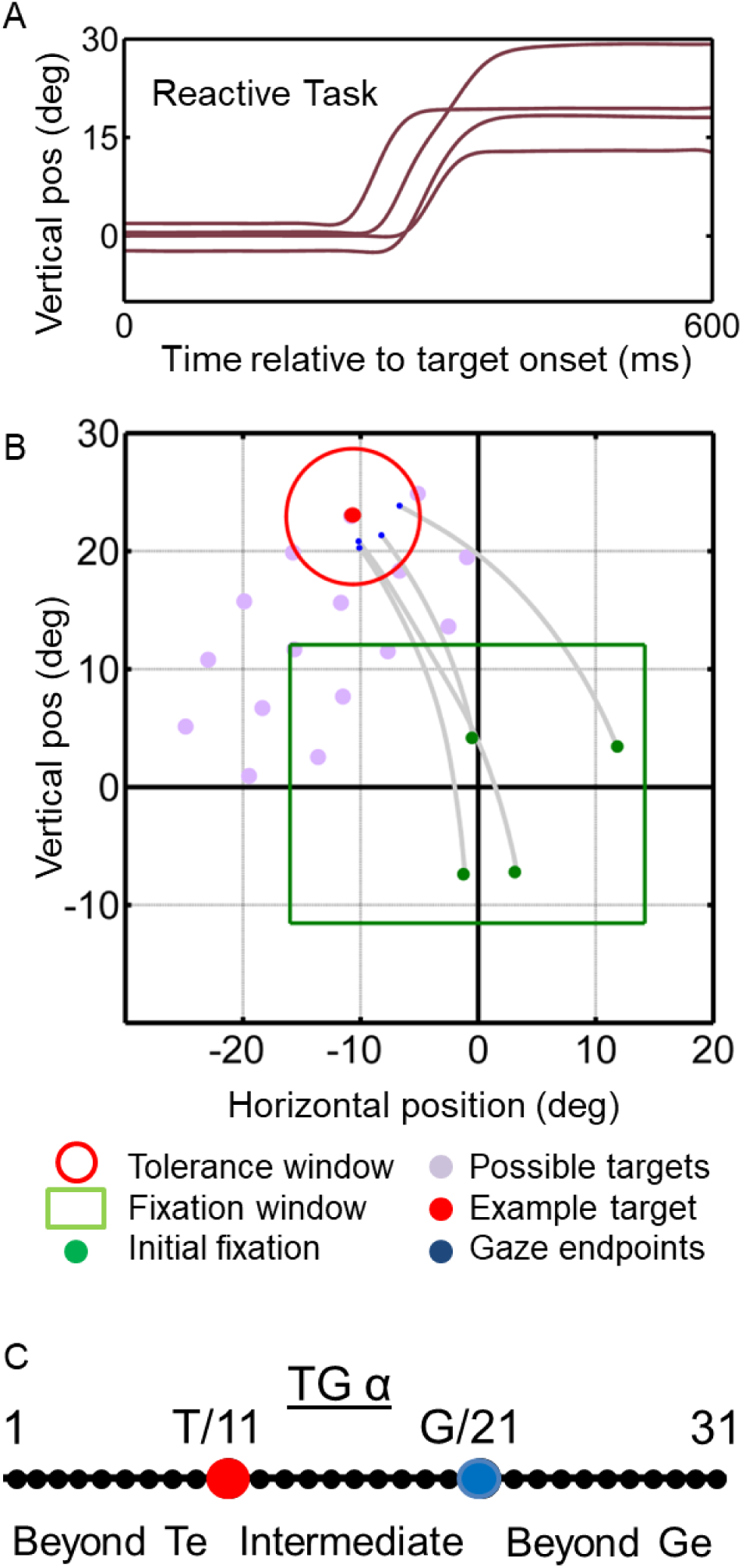
*Reactive gaze task* used for mapping neural receptive fields and fitting models. A) example traces of vertical eye position plotted as a function of time. B) Two-dimensional gaze trajectories (grey lines) from the *reactive task* for an example target in monkey M2. Also shown are the range of initial fixation positions (green square), the tolerance window (red circle), and the other possible targets used in this experimental session (grey circles) to map a neuron’s receptive field. C) The schematic illustrating the target gaze continuum concept, the distance between and beyond the target location and gaze are divided into 31 points and the fit to neural activity is perform at each of the discrete locations to identify the best fit

In addition, we recorded the same neurons in a *Memory delay* task (Sadeh, Sajad et al. 2015). This was identical to the reactive task, except with a memory delay of 400-700ms during which the animal had to maintain fixation before making a saccade. These results were analyzed previously (Sadeh, Sajad et al. 2015) and are only used here to distinguish different neuron types. A more detailed description of eye-head kinematics in this task was described previously (Sadeh et al. 2015); here we focused on gaze kinematics relative to target location.

### Trial definition and inclusion criteria

The beginning of a trial was marked by the appearance of the initial fixation point. The beginning of the gaze saccade was defined as the instant when its velocity exceeded 50°/s, and its end when its velocity decreased to 30°/s. The contribution of the head movement to gaze is defined here as the head movement from the start to the end of the gaze saccade. However, the head movement was often prolonged after the saccadic component of the gaze shift. Head movements were marked from the start of gaze movement until the point at which the head velocity decreased to below 15°/s. The head movement marks were then visually inspected to ensure correct marks. All trials were considered for analysis irrespective of whether or not the monkey received a reward after the trial. We excluded trials on the basis of spatial and temporal criteria. First, trials in which the directions of the gaze shifts were completely unrelated to the direction of the target (e.g. opposite direction) were removed. Then, we obtained the regression between errors in gaze vs. retinal error (the retinal angle between the fovea and the target at the initial eye position before the gaze shift), and removed trials with gaze error two standard deviations greater than this regression line. Furthermore, every trial was visually inspected, and any trial in which the gaze shift was anticipated (reaction time of < 100 ms after the go signal) and when the gaze shift consisted of multistep saccades was excluded. Finally, for each neuron, we required successful performance for at least 80% of total trials [mean standard error of the mean (SEM) trials = 178(16)], and at least seven successful gaze shifts towards each target location (with a possible maximum of 15, after excluding erroneous trials); also, the neuron had to remain isolated throughout the recording session.

### Neural recordings

We recorded extracellular activity from the left and right SC with tungsten microelectrodes (FHC). The electrode was inserted through a guide tube, which was controlled by a hydraulic microdrive (MO-90S; Narishige International, East Meadow, NY, USA). Isolated signals were amplified, filtered and stored for off-line sorting with the Plexon MAP system. The SC was identified according to criteria published previously (DeSouza, Keith et al. 2011, Sadeh, Sajad et al. 2015). The steps of SC identification and confirmation are identical to those explained previously (Sadeh, Sajad et al. 2015). The memory delay saccade task was used to dissociate between visual and movement related activities and categorize cells into visual, visuomotor (VM) and motor neurons. Visual neurons were defined as cells that showed a robust burst of activity (> 50 spikes/s above the baseline) 40–60 ms after the stimulus presentation that lasted for ^~^180 ms afterwards (Goldberg and Wurtz 1972). Motor neurons were those with robust activity or a buildup of activity peaking at the time of gaze onset, with activity starting prior to the gaze onset (100–40 ms before saccade), and that continued to ^~^100 ms after gaze onset. Neurons that met both criteria were classified as visuomotor. We also used a visuomotor index (VMI = (Motor spike count - Visual spike count / (motor spike count + visual spike count) to quantatively separate these based on our previously published memory-delay task data (Sadeh, Sajad et al. 2015). The visual and motor burst spike counts were first subtracted from the baseline activity (100ms pretarget period). This gave a score where −1 is a purely visual neuron and +1 a purely motor neuron). Neurons classified as visual had VMI values ranging from −0.83 to 0.42, visuomotor neurons ranged from −0.74 to 0.51 and the pure motor neurons had VMI values from −0.2 to 0.74. When we refer to ‘number of spikes’ below, this refers to number of action potentials in these defined temporal windows, also we use neural activity and burst interchangeably to refer to the same concept of high frequency of action potentials.

The temporal windows that we used for analysis of bursting activity in the reactive task are illustrated in the results section (Figure 2). For some analyses (i.e., Figures 3, 4) we used a fixed window of +70 to +170ms relative to visual target presentation for visual activity (shown as red vertical lines) and −50 to +50 ms relative to saccade onset (shown as black vertical lines). For other analyses (i.e. Fig. 5, 6) we considered the entire burst duration of the neurons (windows shown as blue vertical lines). The average range of the entire population burst (aligned on stimulus presentation) was 342 ms. For visual neurons the full duration of burst was defined as the time which the activity increases above 50 spikes/s after the stimulus presentation to a point detected by visual inspection at which the activity considerably declines, this window was on average from +48 ms (start) to +231 ms (end) relative to visual stimulus onset. For VM neurons the average range of the entire burst was +47ms to +421 ms relative to visual stimulus onset. For motor neurons the average range was −94 to 194 ms relative to saccade onset. Finally, for figures 6 and 7, we performed a step wise analysis of the entire duration of individual neuron activities broken down into smaller time windows in order to investigate changes in spatial coding during the neural activity (see ‘spatiotemporal analysis’ approach below).

**Figure 2:**
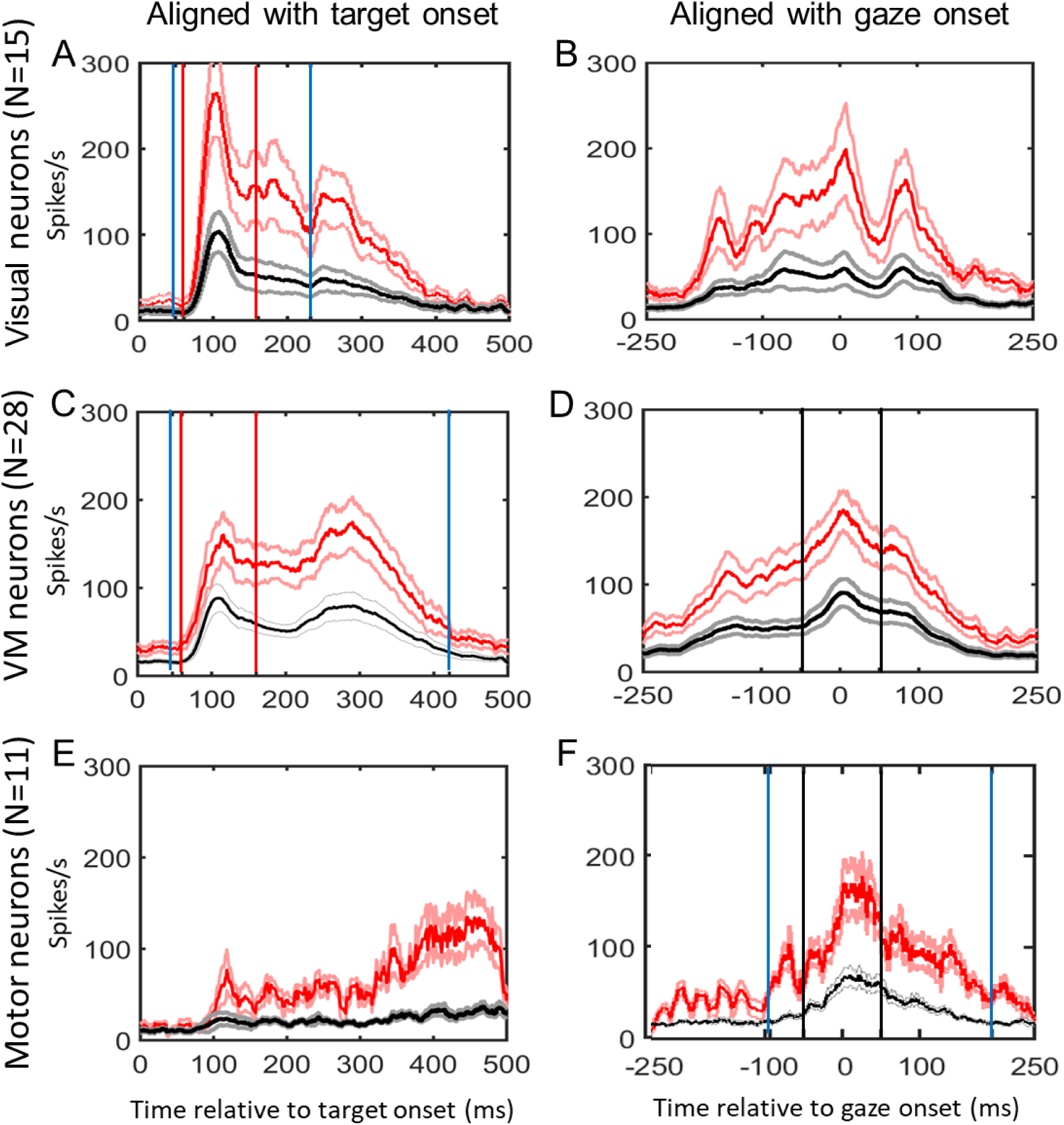
Mean spike density plots for our visual neurons (A and B, N=15), Visuomotor neurons (C and D, N=28), and Motor neurons (E and F) N=11) during the *reactive task*. These subpopulations were identified using the *memory delay task* (Sadeh et al. 2016), not shown here. Data are aligned with stimulus onset (*left column*) and gaze movement (*right column*). These data were averaged across all data that passed our exclusion criteria. Red lines represent spike density plots derived from the ‘top 10%’ trials in the reactive task (±SEM, light red lines), generally corresponding to the RF ‘hot spot’, and the black lines are derived from the average firing rate across all trials (±SEM, grey lines). Solid blue vertical lines indicate the average temporal analysis window for the ‘full burst’ analysis, whereas red and black vertical lines indicate the time intervals sued for the ‘fixed window’ analysis in visual and motor activities respectively.

**Figure 3:**
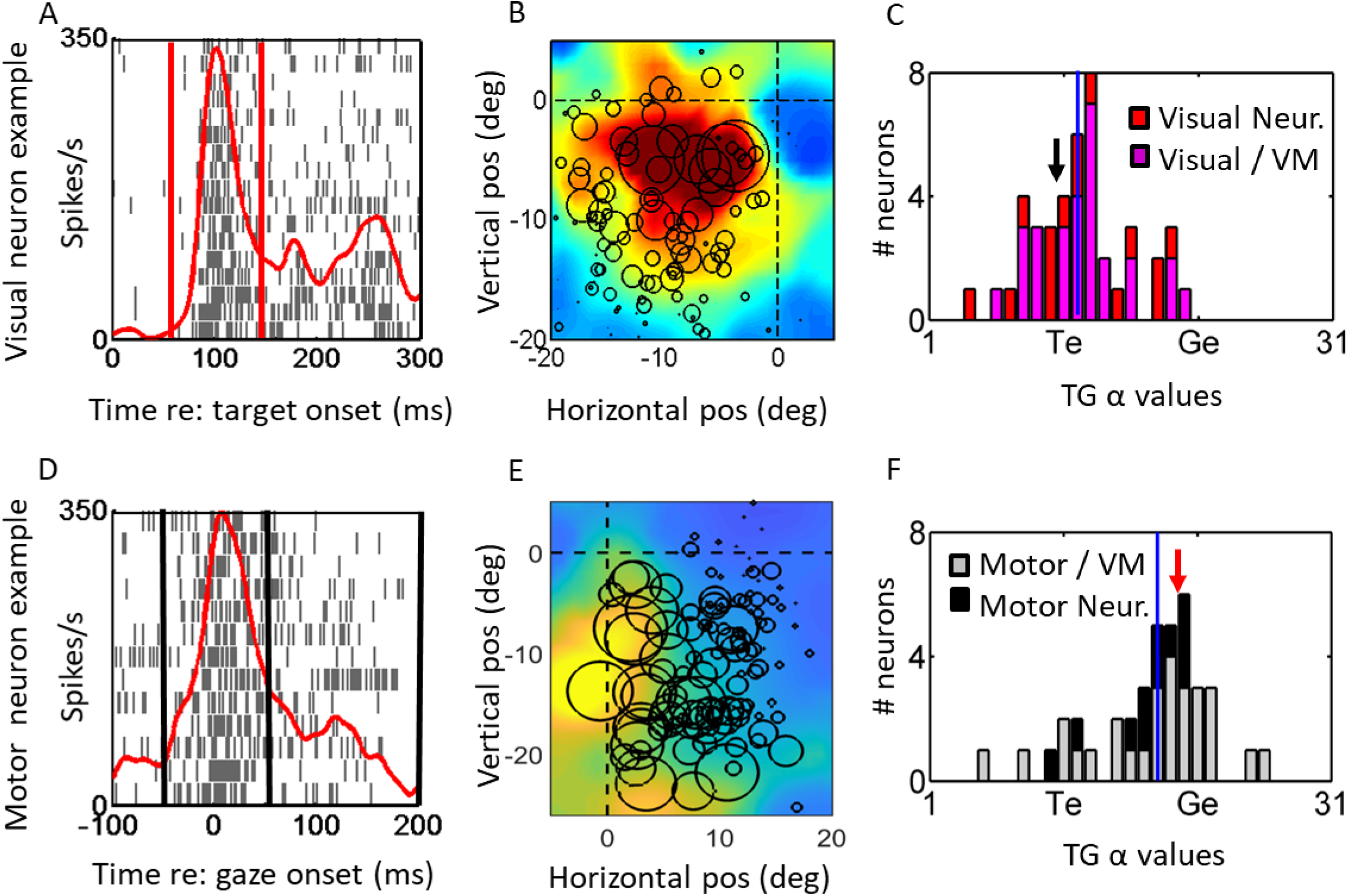
Shift of spatial representation from near Te in the target-aligned fixed window analysis (*top row*, A-C) toward Ge in saccade-aligned fixed window analysis (bottom row, D-E) of *reactive task* data. Each row shows the raster / spike density plot (*left column*) and best fit response field (middle column) for an example neuron, followed the distribution of TG alpha values of full population. For the visual response population (C) both visual (red bars) and visual activity of VM neurons (pink bars) are included. For the motor response population (F) the motor activity of VM neurons (grey bars) and motor neurons (black bars) are shown. The red/black vertical lines in the raster plots (A/D) represent the fixed visual/motor temporal windows respectively. The black vertical lines in the histogram plots (C/F) represent the median TG alpha values and the location of TG value for the representativeexample is indicated by the red arrow. The cluster of the distribution of visual fits (C) is closer to Te whereas the cluster of motor fits (F) is closer to Ge. Note that the shift from the mean TG values in the visual activity histogram (C) (mean= 12.2) is significantly different (unpaired two tailed t-test, p=0.0001) from the mean in the motor activity TG histogram (F) mean (= 17.4)

**Figure 4:**
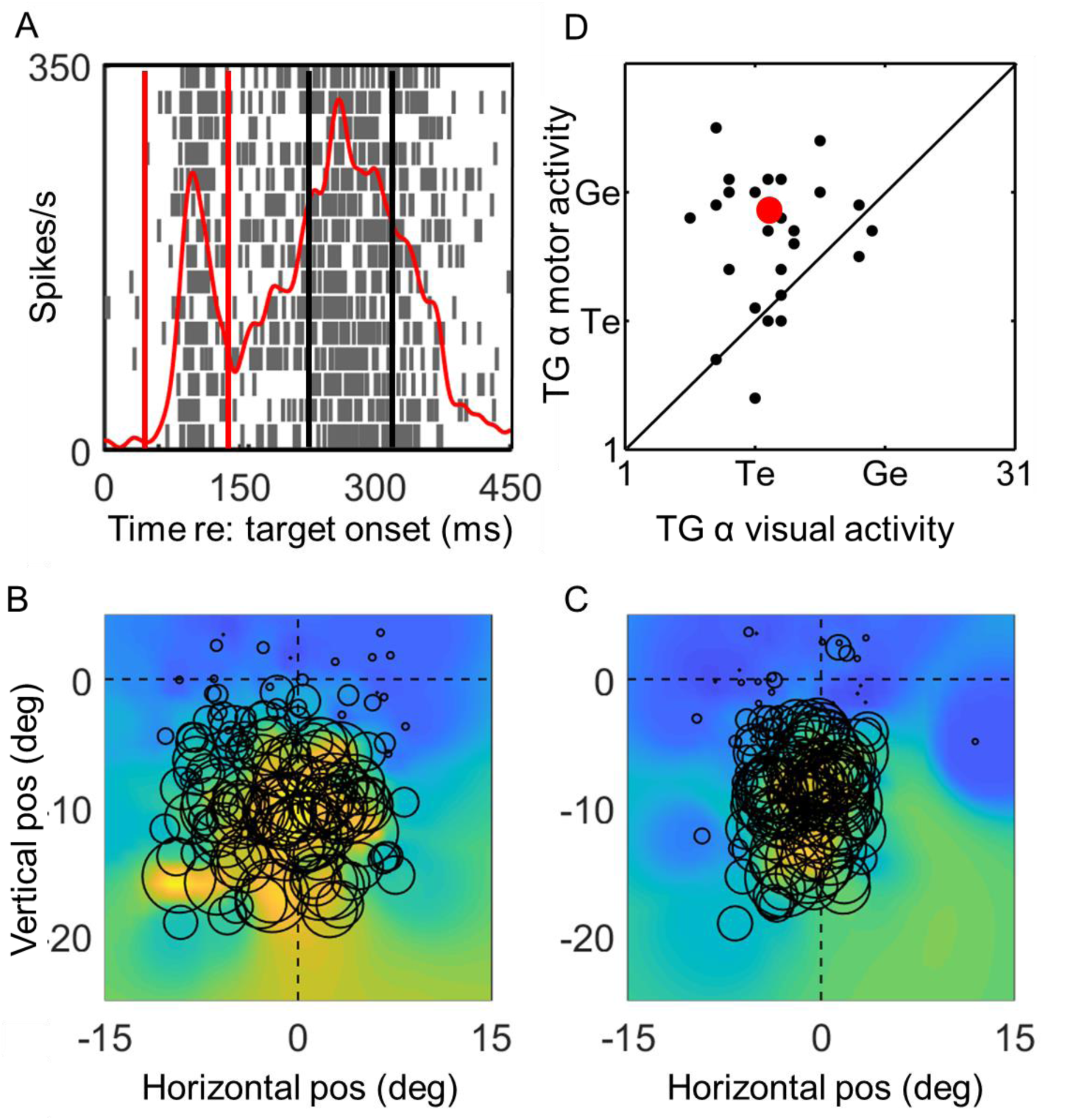
Shift from Te to Ge coding *within* VM Neurons. A) Raster/ spike density plot of a representative VM neuron aligned on target onset, showing fixed visual window (red lines) and average location of fixed motor window (black lines). This is followed by the best RF fit plots for the fixed B) visual and C) fixed motor activities. D) The scatter plot of differences in TG alpha values of visual (x axis) and motor (y axis) of visuomotor neurons (black circles) relative to the equality diagonal line. The average of the TG alpha values in represented by the red circle and the representative example shown in 4A-C is indicated as the red circle. Most neurons lie above the line which indicates that there is a transition from coding for target location in the visual activity to gaze end location in the motor activity within the individual VM neurons. This shift was significant (paired two tailed t test, p=0.001).

**Figure 5, A-I:**
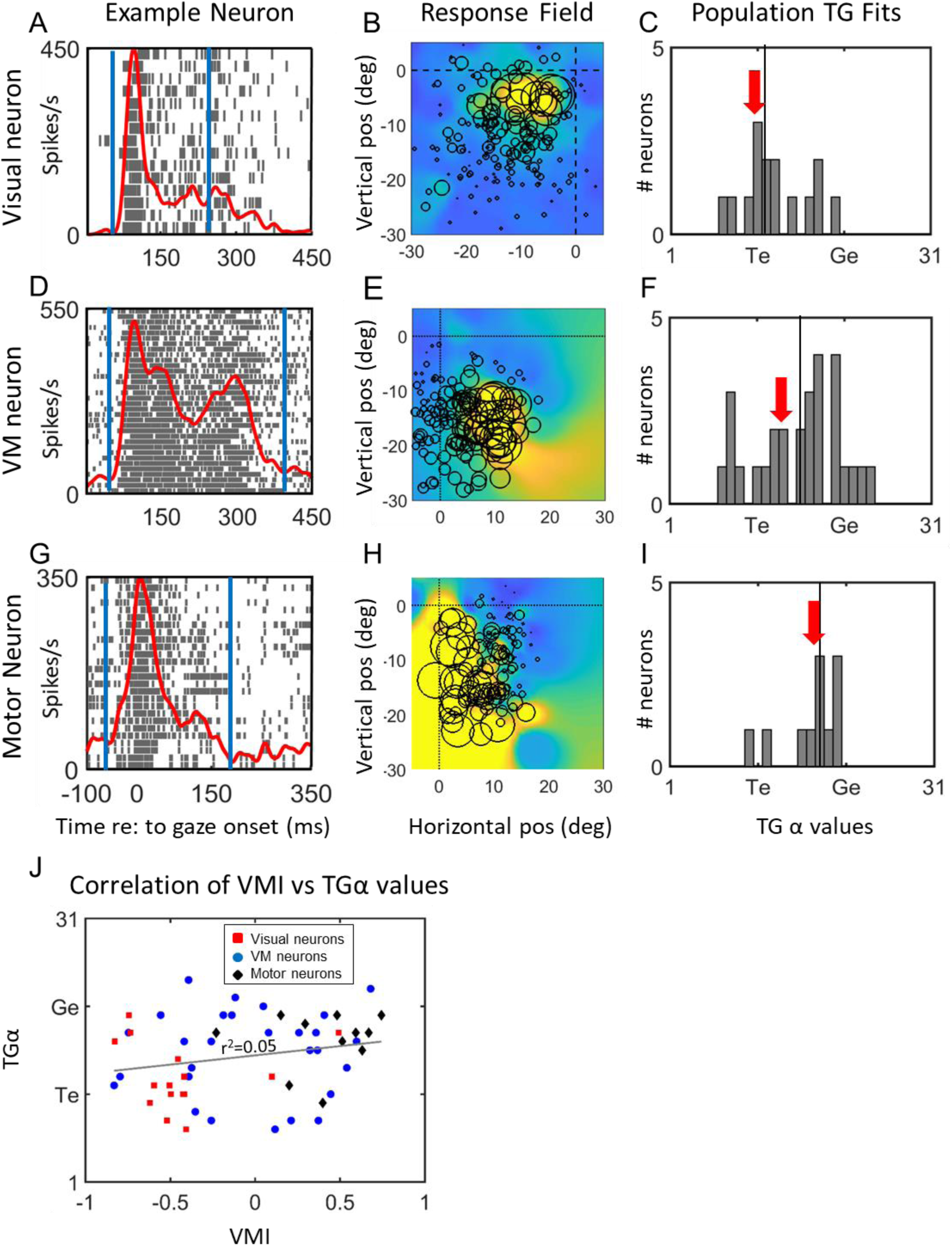
The TG alpha value distribution for ‘full burst’ analysis of visual (top row 1, A-C), VM (row 2, D-F) and motor (row 3, G-I) neural populations in the reactive task. Each row shows an example neuron’s spike density plot/raster (column 1) and receptive field (column 2), and then a frequency histograph of best TG fits for all neurons in that population. Spatial fits were made for each neuron using data derived the entire duration of task-related neural activity (between blue vertical lines in *left column*), aligned on stimulus onset. The vertical line in each panel of the *right column* indicates the median of the TG alpha values and the red arrow indicate the TG value of the representative example. J) TG alpha values plotted as a function of visuomotor index (VMI) for each neuron population. All neuron categories exhibit a weak, non-significant correlation: Visual neurons are represented by red squares (r2=0.0119, p=0.7), VM neurons by blue circles (r^2^=0.0012, p=0.86) and motor neurons by black diamonds (r^2^=0.001, p=0.98). The overall correlation across all neurons (indicated by the gray correlation line) also leads a weak (r^2^=0.05) non-significant(p=0.1) correlation between the two variables.

**Figure 6:**
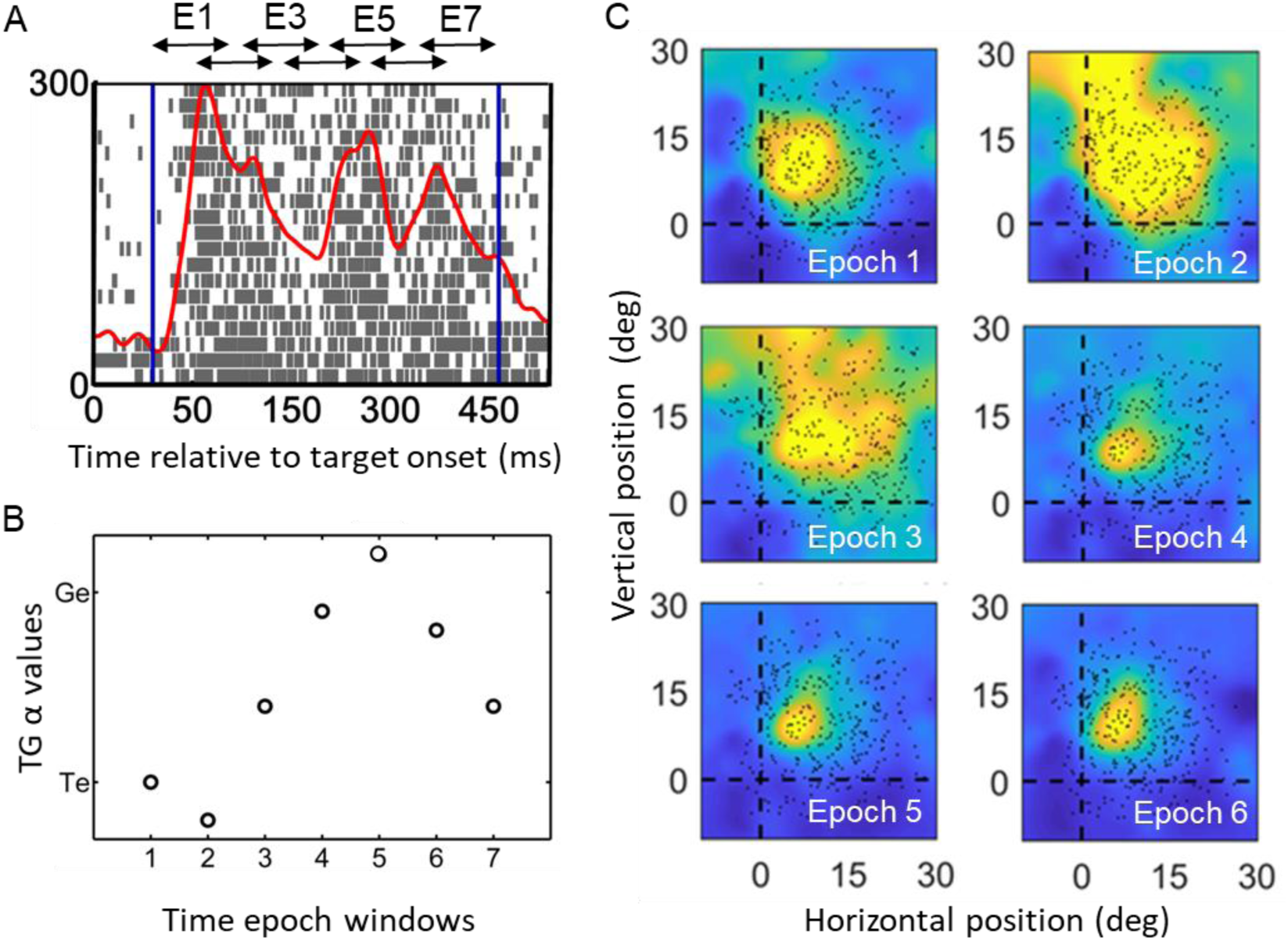
Spatiotemporal analysis in one example neuron. A) Action potential raster plot and spike density plot of a representative visuomotor neuron during the *reactive task*. The spike density plot (thick red line) was derived from the trials with the top 10% of activity (N=19), i.e., when the target was presented at the ‘hot spot’ of the RF. The dark blue vertical lines indicate the sampling window of the entire visuomotor burst. The double headed arrows on top of the raster plot indicate the semi-overlapping time windows which were used for the response filed and TG value analysis shown in B and C. These sampling windows were normalized according to the duration of the action potential (-370 to 200 ms relative to gaze onset) to yield 7 overlapping windows with equal time periods. B: TG continuum values plotted as a function of their sequence through time (1-7). In this case there is a rise from T toward G over the first 5 steps followed by a slight reversal. The details of these patterns varied across neurons. C: RF fits for the activity from time windows 1-6-, plotted in the best fit reference frame along the Target-Gaze Continuum (epoch 7 looked the same as 6). The dots indicate spatial positions of the targets in this frame for each trial and the color heat map (blue = low activity, red = high activity).

**Figure 7:**
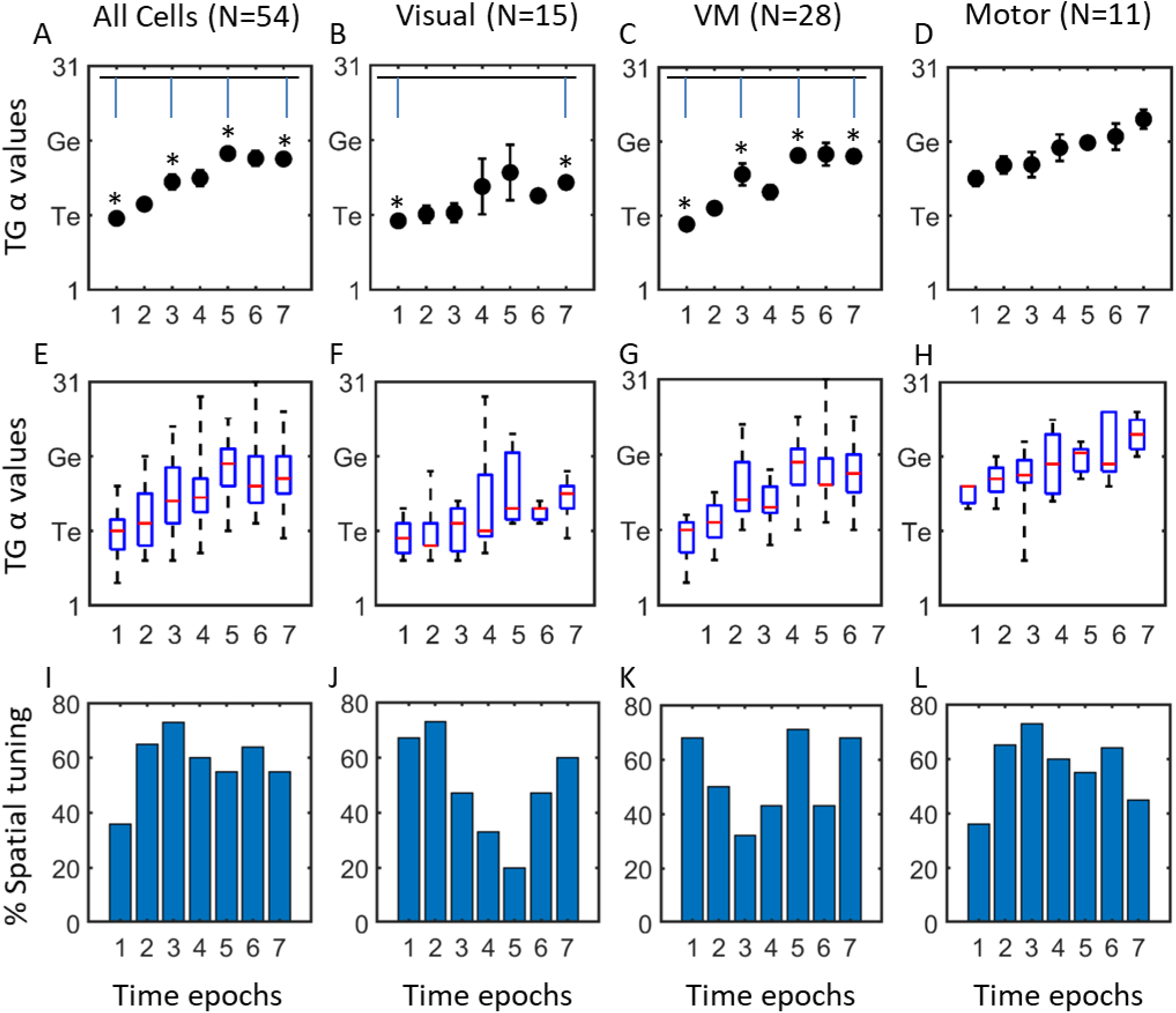
Spatiotemporal analysis in entire superior colliculus neuron population (*column 1*) and each sub-population (*columns 2*-*4*). *Top row* (A-D) shows the mean TG alpha values (y axis) of each temporal window of analysis (x axis) with SEM bars, the *middle row* (E–H) shows the median values (red bars) as well as first and third quartiles (blue bars) of TG alpha values (y axes) for the same data, and the *bottom row* (I-L) shows the percentage of cells in each time epoch that showed significant spatial tuning. The entire neuron population (*Column 1*, N=56), showed a gradual shift in each step from more Te related coding in the earlier visual activity to more Ge related as the activity becomes closer to gaze onset. The Visual neuron population (*Row2*, N=15) which showed a predominantly preference in coding for target especially in earlier windows with a non-significant shift toward intermediate TG alpha value later in its activity (one-way ANOVA p=0.402). The VM population (*Row 3*, N=28) showed a significant shift in TG alpha values (One-way ANOVA p=0.0001). The Motor population (Row 4, N=11) started at a more intermediate TG value and showed a non-significant shift toward G (one-way ANOVA p=0.48). The significant differences (P<0.05) are indicated by asterisk (^∗^). However, as described in the text, there was no significant difference between these three patterns. Note that for the results shown in Fig 5A-H, the TG values were included in the analysis only if the neuronal activity showed spatial tuning for that given analysis window.

### Spatial Analysis of Neuronal Response Fields

Visual and motor RFs were obtained for each neuron for all of the models and in order to analyze and compare the spatial coding we used several spatial models to fit the RF data for each neuron using a method that has previously been described several times (Keith, DeSouza et al. 2009, DeSouza, Keith et al. 2011, Sadeh, Sajad et al. 2015, Sajad, Sadeh et al. 2015). Briefly, the RF of the neuron was plotted by overlapping firing rate data over two-dimensional position data corresponding to the spatial parameter related to the given model (e.g., final gaze position relative to the eye; for the list of models tested in this study see below). Spatial models were then constructed by fitting the RF data non-parametrically using Gaussian kernels with bandwidths ranging from 2-15 degrees. The qualities of the model fits were quantified by calculating the Predicted Sum of Squares (PRESS) residuals for all trials, which is a type of cross validation in regression analysis (Keith, DeSouza et al. 2009). The spatial code of a neuron was then defined as the model (at the kernel bandwidth) that yielded the overall best fit (i.e. smallest residual) to the data. Briefly, PRESS residual for every trial was obtained by: 1) eliminating that trial from RF data, 2) fitting the remaining data points non-parametrically using Gaussian kernels at various bandwidths (2-15°), and 3) obtaining the residual between the fit and the missing data point. The overall predictability power of the model for the recorded data set was quantified by the average of PRESS residuals across all trials for that neuron.

As noted above, the spatial parameters in our behavioral task (Figure 1) were designed to distinguish between various frames of reference using the analysis described above. These were tested exhaustively in a previous analysis of neurons recorded in the memory delay paradigm (Sadeh, Sajad et al. 2015); (which used an overlapping but larger population of neurons) we tested eleven models that have been proposed for spatial coding in the eye and head movement control system against the visual and movement responses of all neurons. This included models of target location vs. gaze, eye-in-head, and head motion (both final position and displacement) in eye-centered, head-centered, and body-centered frames of reference). This yielded an overall preference of SC neurons for target (T) and gaze (G) position codes described in eye-centered coordinates. These results allowed us to narrow down and refine our spatial models to examine neuronal coding along a continuum of intermediate spatial models spanning T and G.

The physical basis of the TG continuum is illustrated in Figure 1 C, which shows the TG continuum for an example trial. This continuum extends between, and beyond T and G position for every such trial. The intermediate spatial models were constructed by dividing the distance between target position and final gaze position for each trial into 10 equal intervals and 10 additional intervals extended on either end. The location of the best-fit model along the T-G continuum (here referred to as TG alpha value) is indicated by a value between 1 to 31 (the Target and Gaze locations are arbitrarily numbered 11 and 21 respectively) indicating their relative preference for coding target vs. gaze related spatial information. For example, if the fit and TG continuum analysis for the activity of a given neuron yields the value of 20 (one step away from 21 – i.e., G), this indicates that the spatial information encoded by this neuron’s activity is best described by a position between target position and gaze endpoint that’s 90% described by gaze endpoint, and only 10% by target position. Noteworthy that this analysis is not influenced by systematic errors in behaviour and entirely relies on variability in the spatial relationship between positions in different models. Once the optimal TG value is determined, it can then be used to plot neural RF’s in their intrinsic coordinate system, simply by plotting activity for trial according to its location along the TG continuum (in eye-centered coordinates).

### Spatiotemporal Analysis

In order to track changes in the spatial code through time (Figs 6, 7), we used a step by step analysis of the entire duration of the burst when broken down into smaller time windows, i.e. analyzing each time window separately using the same model fitting approach. The specifics of the analysis approach were explained in detail previously (Sajad, Sadeh et al. 2016), but briefly: the similar spatial analysis as described above was applied to each of the time windows spanning the visual and motor neural activities and in order to account for variabilities in duration of the activities from one neuron to another without losing any of the activity in analysis we normalized the time between the onset of modulation aligned on target onset based on spike density function (mean = 57 ms after target onset for V and VM neurons, and 86 ms for motor neurons) and the time of gaze movement onset which varied on a trial by trial basis for all trials, the duration between this early visual period and gaze movement onset was on average 231 ms (± 74 ms, SD) across all trials. The normalization served to account for time and space similarly since the T-G continuum data are also obtained by dividing the spatial difference between target position and final gaze position (i.e., inaccuracy errors in behaviour) in fixed number of discrete steps on a trial by trial basis The analysis on the RF sampled from the activity within the time-normalized windows allows for the visual and motor activities to be analyzed as a continuum to detect possible gradual changes in spatial coding through time.

The firing rate of the neuron in the corresponding window (spikes/sec; number of spikes divided by the sampling interval for each trial) was sampled at 7 semi-overlapping windows from this time-normalized data. This choice of sampling window numbers was based on the approximate ratio of the duration of the visual response to decrease in the peak and then to the start of movement response including a post-saccadic period starting from gaze onset. The final (7th) time-step corresponded to mostly post-saccadic period starting from the onset of gaze shift. Because of the time-normalization process the sampling window width scaled with the duration between visual response onset and movement onset on a trial-by-trial basis. On the 7-step time-normalized scale, the visual burst on average lasted 4 steps (SD = 0.63 steps), ending by the end of the fourth time-step in 91.2 % of trials. The sampling window width was on average 75ms (±8ms, SD) and was no less than 47ms for any trial which ensured enough neuronal spikes captured in the sampling window to perform effective spatial analysis. The time for which the first window starts was also confirmed by visual inspection of activity raster of all neurons to identify the visual bursts, movement bursts and the peaks.

### Confirmation of significant spatial tuning (in neuron populations)

Since the results of our analysis approach are only considered valid if the sampled neural activity exhibits spatial tuning, we excluded any data point which did not exhibit significant spatial tuning. In order to achieve this, we used an approached described in details before (Sajad, Sadeh et al. 2016) randomly shuffled the neural activity data and plotted the data over the positional data of the best fit model for the neuron to obtain a ‘random’ RF. This process was repeated 100 times and therefore 100 random RFs were obtained. To do this, we randomly shuffled the firing rate data (number of spikes divided by duration of the sampling window) and plotted them over the position data corresponding to the best-fit model, and repeated this procedure 100 times to obtain 100 *random* RFs. The PRESS residuals of these random RFs (and their respective mean PRESS values) were then obtained after fitting the data (non-parametrically, using Gaussian kernels) with the same kernel bandwidth that was used to fit the best-fit model, resulting in a total of 100 mean PRESS residuals. If the mean PRESS residuals for the best-fit model (PRESS _best-fit_) were at least 2SD smaller than the mean of the distribution of random mean PRESS residuals, then the sampled activity was categorized as spatially-selective. Moreover, in order to exclude any non-spatially tuned activity and reduce the overall noise to signal ratio in our population we excluded population data belonging to time-windows at which the mean spatial coherence of the population was not significantly higher from that of the baseline activity prior to target presentation which demonstrates no spatial tuning. We used a coherence index (1 - (PRESS_best-fit_/PRESS_random_) value in order to determine the contribution of each neuron to the overall spatial coherence of the population (Sajad, Sadeh et al. 2016).

## Results

### General Observations

We sampled 86 SC neurons during head unrestrained gaze shifts. Of these 86, we were able to record a complete data set from 74 neurons, spanning both sides of the SC in each animal. Of these 74 neurons, 54 met all of our inclusion criteria, including 15 visual, 28 VM and 11 motor neurons (as Identified using the memory delay task; (Sadeh, Sajad et al. 2015)).

Figure 2 shows the activity profiles of each category of neurons (Visual, VM, Motor) during reactive gaze saccades to the top 10% RF ‘hot spot’ (i.e. the region of the RF with the highest neural activity) data (red traces) and the full RF dataset (black traces). Each panel provides mean spike density plots (averaged across neurons ± SEM). Data are aligned both with target onset (Left column; Fig 2 A, C and E) and when aligned with gaze onset (Right Column, Fig 2B, D and F). Vertical red and black lines indicate the ‘fixed-window’ visual and motor analysis windows respectively, whereas blue vertical lines indicate the average duration of the ‘full burst analysis’. (Note that Figure 2 shows average full burst durations for neuron populations; some neurons burst for shorter or longer durations but sum over the whole range, so the mean population spike density plots show a longer duration than the mean full burst windows).

By definition, visual neurons showed a much stronger target-aligned response than saccade-aligned response (Fig. 2 A vs. B), VM cells showed approximately equal responses (Fig. 3 C vs. D), and motor neurons showed much stronger saccade-aligned responses (Fig. 3 E vs. F).

The visual neuron population showed a strong initial peak of activity 48 ± 11 ms (mean ± SD) after the stimulus onset, followed by a smaller secondary peak of activity at 210 ± 15 SD ms (Figure 2 A). The large third peak 300 ms past stimulus onset was likely residual motor activity (i.e., not excluded by our *memory saccade*-based population criteria) because it was absent in the memory delay task visual response (Sadeh, Sajad et al. 2015), and aligned closely with saccade onset (Figure 2 C). This was excluded from the visual full burst analysis, except in the stepwise temporal analysis shown below (Figs. 6,7).

The VM population showed a first peak 106±9 ms after the visual stimulus onset (Figure 3-2B) and a second peak 9±3 ms after saccade onset (Figure 2D), separated by a short period (average 95 ± 12 SD ms) of sustained activity. Motor neurons showed a single peak of activity 22 ± 6 ms) after saccade onset (Figure 2F). Henceforth we will refer to the data from our fixed target and fixed saccade-related windows as ‘visual activity’ and ‘motor activity’, based on their temporal profiles, but use our TG-continuum analysis method to quantify what spatial parameters these activities actually encode in different neurons and at different times.

### Spatial Transformation between Visual and Motor Responses

In our previous papers (Sadeh, Sajad et al. 2015) we used fixed visual and motor window analysis in combination with a memory delay paradigm to show that SC and FEF visual responses tend to code Te whereas the motor responses, following a brief memory period, tends to code Ge. The spatiotemporal analysis described above suggests that the same is true during reactive saccades, i.e., even in the absence of a memory delay. To test this directly, we repeated a fixed visual/motor window analysis on the reactive task data (see Methods and Figure 2). Note that these two temporal windows were each 100 ms in duration, and on average were shifted from each other (start-to-start) by 192 ± 23 ms, meaning that they were separated end-to-start by only 92 ± 23 ms. Thus, we were testing if a significant spatial transformation from T toward G coding occurred over a very short period of time.

Figure 3 provides example rasters and fixed analysis windows (left column) and RF fits (middle column) for a typical visual cell (top row; A, B) and motor cell (bottom row; C, D). The right column provides frequency histograms and scatter plots that contrast the TG alpha values for visual and motor window fits for our entire population of cells. The results of the visual window analysis are shown in Fig 3C. Overall, this yields a mean (12.2) and median (12) and distribution (SD 4.2) that clearly clustered near Te (11). There was no significant difference between the mean of the means of TG alpha values for the visual population (red bars) and the visual response of the VM neurons within the same time window (pink bars) (p= 0.8738, unpaired t-test). In contrast, our analysis of motor activity (Fig. 3 B) yielded an overall mean (17.3), median (18), and distribution (SD 4.7) that was shifted toward the Ge model. Again, there was no significant difference between the distribution of the motor neuron responses (black bars) versus the motor response of VM neurons (gray) within the same time window; (unpaired t-test, p=0.85. More importantly, there was a significant difference between the distributions of the visual (Fig 3A) and motor (Fig. 3B) responses (P= 0.0001, unpaired t-test)

Remarkably, this rapid shift in coding can be observed even *within* VM neurons, such as the example neuron with raster / spike density plot shown in Figure 4 A, visual receptive field in Figure 4 B, and motor response field 4 C. To directly quantify if a TG shift occurs *within* VM neurons, we plotted the TG alpha value from the motor window as a function of the value of the visual window for each neuron (Fig. 4 C). Neurons with data points that lie above the diagonal line indicate a different preference of spatial coding in their visual versus movement related activities. The mean of TG values for VM neurons is also indicated by a red circle in Figure 4D which shows that as a population there is a shift from target to gaze coding when going from visual to movement related activities in the VM neurons. Overall, the motor TG values for VM neurons were significantly different from their visual TG values (Paired t test, p= 0.0001). Thus, a rapid transformation along the TG continuum occurred between visual and motor responses, even within VM neurons.

This analysis suggests that the spatial code in SC neurons is not stable during a reactive task, particularly within VM neurons. However, it is not yet clear to what degree the overall visual motor transformation is influenced by the spatial contributions of different neuron types at different times. This is not trivial to answer, given that visual cells by definition are active before motor cells, this classification scheme and timing will interact. Does this visuomotor transformation occur because 1) neurons with early responses have a fixed T code whereas later motor neurons show a fixed G code, 2) because a distributed transformation causes a spatial shift in the code of late responses away from T, or 3) due to some combination of these factors? The first possibility (cell-fixed coding) does not seem compatible with our VM data (Figure 4D), but we performed a more in-depth analysis explore this in more detail.

### TG Continuum in the full burst of visual, VM, and motor cell

To test whether there is an overall difference in spatial coding between our three different neuron types (V, VM, M) could be influenced by a fixed neural code in each cell type, we analyzed the full burst (Figure 2) of each neuron types. In a previous paper (DeSouza, Keith et al. 2011) a similar model-fitting approach was used on the full burst of Superior Colliculus neurons during the reactive task, but that study did not use a memory-delay task to classify different neuron types, and did not provide a TG continuum analysis (only ‘cardinal’ models such as Te, Ge, etc.). Based on that analysis DeSouza et al. (2011) concluded that the Superior Colliculus burst primarily encodes Te, but the current analysis provides a more nuanced picture.

Figure 5 shows the ‘full burst analysis’ for our visual neurons (A-C), VM neurons (D-F) and motor neurons (G-I) respectively, showing an example neuron (left column), its RF at the TG value of best fit (middle column), and the frequency distribution of TG-α for each population (right column). The entire combined population (not shown) generated a TG alpha median of 16.5 (SD=4.4), roughly in the middle of the T-G continuum (TG-alpha = 16). However, the distribution of individual neuron fits was quite broad and possibly clustered near T and G, perhaps suggesting the co-existence of different spatial codes. When these data were divided into different types, however, visual neurons (Fig. 5C) were clustered toward Te (11), with a mean TG score of 13 (SD=3.8), VM neurons (Fig. 5F) continued to show a broad distribution, with mean of 15.8 (SD= 4.9), and motor neurons (Fig. 5 I) clustered toward G (21) (mean: 17.9, SD=3.3). This analysis shows 1) that superior colliculus neurons show a broad continuum of spatial tuning between T and Ge during the reactive task, and 2) that different neuron types made a slight, significant (One-Way ANOVA, p=0.04) different contributions to this distribution, with visual cells clustering toward Te, Motor cells clustering toward Ge, and the distribution of VM cells spanning both.

Despite these tendencies, each sub-population showed a distribution of fits along the TG Continuum (Figure 5 C, G, I). To test if this was due to variations in Visual-Motor tuning within cell types, we correlated the TG fit of these cells obtained from their full burst in the reactive task against their visuomotor index (VMI) obtained from the same cells in our memory delay task (Sadeh, Sajad et al. 2015). The overall relationship is shown in Figure 5J, with each sub population coded for color. This yielded very weak correlations for visual (r^2^=0.0119, p=0.7), visuomotor (r^2^=0.0012, p=0.86) and motor cells (r^2^=0.001, p=0.98). Even the entire cell population only showed little correlation between TG score and VMI (r^2^ = 0.05, p = 0.1), suggesting that the relative size of the visual vs. motor burst was not the main determining factor in the spatial codes in these cells.

### Spatiotemporal progression of visuomotor Signals in the SC

To test if timing is the key factor in determining the spatial code in SC cells during our task, we examined the progression of spatial code through time for each neuron. Specifically, the entire activity of each of the individual neurons in each category was divided into seven time windows using a time normalization method to account for differences in duration of activity (See methods), the resultant TG alpha value was combined for each individual window in each of the neuron categories in order to investigate the temporal progression and transformation of spatial codes in each of the populations (Sajad, Sadeh et al. 2016).

Figure 6 illustrates this analysis using an example VM neuron. Figure 6A illustrates that this neuron had multiple peaks of activity, including an initial visual peak, a strong secondary visual response, and a motor response. Figure 6B shows the corresponding RFs of the first 6 windows (each plotted using is optimal fit on the TG continuum), showing how they progress through time. Figure 6 C then shows these TG fits as a function of time. Note that although these fits often ‘bounce around’ for individual neurons like this example, especially near the start and end where spike rate is rising and dropping and confidence is thus lowest, they show a general trend to progress from near T to nearer G, as one can see in the next analysis.

To test the temporal shift in spatial coding at the population level, we first pooled all visual, VM, and Motor cells, and looked at their progression of TG coding across the 388±53 ms duration of their response (Figure 8, first column). Most neurons showed significant spatial tuning during most time steps (bottom row), and only these were used in the TG calculation. Figure 7 A and B demonstrate the mean and median values with SD and SEM bars respectively for each of our 7 normalized time windows, and Figure C shows the percentage of data that was spatially tuned in each window (and thus included in the analysis). The trend of these results suggests a gradual progression of target related coding indicated by TG values closer to the T model (i.e. TG=11) in earlier more visually related activity to gaze coding (values closer to TG of 21) in the after activities which are temporally correlated with gaze onset. We compared the TG values in time windows 1, 3, 5 and 7 to exclude comparison between the overlapping windows using Kruskal–Wallis non-parametric one-way ANOVA test and found an overall significant difference (p<0.0001) between the windows. We also found significant differences in TG value of window 1 (mean: 11.1) compare to value of windows 3, 5 and 7 (means: 14.7, 19.6 and 18 respectively and P<0.01, P<0.001 and P<0.001 respectively). Further, the relationship between TG code and timing of the response yielded a very strong correlation (r^2^=0.94 p<0.00001).

### Timing vs. Cell Type

As noted above, timing and a cell classification based on visual-motor balance could interact or mask each other’s effects. As a result, cell type differences could look like timing differences and vice versa. To disentangle these effects, we divided our time analysis data into separate visual (Fig 8. 2^nd^ column), VM (third column), and motor (fourth column) populations, they each showed similar trends, except that the ‘visual’ population code plateaued before reaching G. Note that over the course of our seven time steps, the percentage of spatially tuned visual cells (shown in the bottom row) peaks around the time of the late visual response and fades toward the saccade, whereas spatially tuned activity held steady in the VM population and ramped up in the motor population. Testing within the three populations, there was a significant difference between first and seventh time steps in the visual neuron population (P=0.03) and between the first and third, fifth and seventh time steps in the VM neuron population (P=0.01, P=0.001 and P=0.0001 respectively). No significant changes in the TG values were observed between the time windows in the motor neuron population, but each population showed a significant correlation as a function of timing: Visual neurons: r^2^=0.6, p=0.0006, VM neurons: r^2^=0.81, p<0.00001, and Motor neurons: r=^2^0.96, p<0.00001.

Based on visual inspection, there appears to be a slight upward shift (from T toward G) in these time-normalized plots from visual (Fig. 8 B), to visuomotor (Fig. 8C), to motor (Fig. 8 D) populations. However, there was no significant difference between these plots (P = 0.53, Nonparametric One-way ANOVA test test). These results suggest that a similar spatiotemporal progression occurs across different cell types in the SC during reactive saccades, and that the difference in spatial coding across different cell types (Figure 3) are primarily due to the relative timing of their responses, rather than fundamental differences in neuron properties.

## Discussion

The process of transforming the visual information into movements command must occur for a successful and timely gaze shift (Mays and Sparks 1980, Gnadt, Bracewell et al. 1991, Crawford and Guitton 1997, Pouget and Snyder 2000, Snyder 2000, Crawford, Henriques et al. 2011, Sajad, Sadeh et al. 2015, Sajad, Sadeh et al. 2016). Here we found that the superior colliculus (SC) participates in a rapid transformation from target to gaze coding, even in the absence of a memory delay or other experimental manipulations. Further, we have shown this does not primarily arise because of some fixed intrinsic code within in different cell types (at least along the visual-visuomotor-motor continuum) but rather because of a continuous temporal progression through all cell types. To our knowledge, this is the first direct demonstration of an internal spatiotemporal transformation during simple reactive saccades.

### The Superior Colliculus Spatial Code

It has been a subject of debate whether the SC codes T, target location (Sparks and Porter 1983, Waitzman, Ma et al. 1988, Sparks 1989, Basso and Wurtz 1998, McPeek and Keller 2004) or G, future gaze location (Walker, Fitzgibbon et al. 1995, Freedman and Sparks 1997, Everling, Dorris et al. 1999, Horwitz and Newsome 1999, Klier, Wang et al. 2001). In a previous study (DeSouza et al. 2011), we concluded that overall SC activity preferred a Te code during reactive saccades. In light of the current study, this was likely due to a mixture of different signals and the use of cardinal T and G models rather than the T-G continuum. The current more sophisticated analysis revealed a continuum of T-G codes across all three cell populations, with a preference for T in V cells, a distribution that equally spanned T and G in VM cells, and a preference for G in M cells. This is generally consistent with our analysis of SC activity in a memory delay task (Sadeh et al. 2016), and makes sense in terms V cells presumably reflecting visual input most closely (Wurtz and Mohler 1976, Wurtz and Albano 1980, Moschovakis, Karabelas et al. 1988), motor cells reflecting output (Sparks and Hartwich-Young 1989, Miyashita and Hikosaka 1996, Sparks 2002), and VM cells reflecting both as well as more complex influences. VM neurons are known to receive a more extensive range of inputs from other brain areas (Wurtz and Albano 1980, Moschovakis, Karabelas et al. 1988, Moschovakis, Karabelas et al. 1988, Sparks 2002), have diverse subtypes(Sparks 1978, Wurtz and Albano 1980, Sparks and Hartwich-Young 1989, Munoz and Wurtz 1995, Munoz and Wurtz 1995) and are suggested to be more involved in cognitive and higher order functions (Everling, Dorris et al. 1999, Horwitz and Newsome 1999, Krauzlis, Liston et al. 2004, Sommer and Wurtz 2004, Krauzlis, Lovejoy et al. 2013, Dash, Yan et al. 2015).

### Evidence for a visual to motor transformation in the superior colliculus

One traditional view of spatial coding in the SC is it codes retinal error information received from retina and striate cortex, and simply relays this to the brainstem (Mohler and Wurtz 1977, Distel and Fries 1982, Fries 1984, Waitzman, Ma et al. 1988, Optican 1995; Sparks 2002; DeSouza et al. 2011). Alternatively, it has been demonstrated that the SC (and other cortical gaze areas) can provide a visual-motor transformation for gaze shifts when the experimental task introduced a temporal or spatial separation between the visual stimuli and movement initiation (Gnadt and Andersen 1988, Everling, Dorris et al. 1999, Everling and Munoz 2000, Munoz and Everling 2004, Sadeh, Sajad et al. 2015, Sajad, Sadeh et al. 2016, Sajad A 2016). However, it has been argued that the separation of visual and motor events required in these studies influences spatial code by changing the cognitive demands on the neural circuit, for example forced encoding the target of location by visual activity and the gaze movement by motor activity in the case of antisaccades, or by introducing memory-related errors in the case of a the memory-delay task (Mays and Sparks 1980, Stanford and Sparks 1994, White, Sparks et al. 1994, Goldman-Rakic 1995, Miller, Erickson et al. 1996, Brown, DeSouza et al. 2004, Hollingworth 2015, Sajad, Sadeh et al. 2016, Sajad A 2016).

The current study utilized a simple behavioral paradigm (reactive gaze saccade made directly to targets with no delay), combined with a sensitive model-fitting approach that can track spatial codes based only on endogenous error in the system. Based on the results of our previous study, which tested a wide array of spatial models in a memory delay task (Sadeh et al. 2015) we focused on two models: Target in eye coordinates (Te) and future gaze position in eye coordinates (Ge), and used ‘TG’ continuum between these models to test the visuomotor transformation. The results were clear, even in the short time span (192 ± 23 ms) between our visual and motor analysis windows there was a significant shift in coding across our entire population from T toward a G code. Given the simplicity of the task these cannot be attributed to exogenous suppression, memory, or top-down transformation signals. Instead, we attribute these errors to a transformation occurring within the sensorimotor circuit. Since the output (Ge) still encodes gaze in retinal coordinates, this remains compatible with the notion that the SC provides a two dimensional command to the brainstem in retinal coordinates (Klier, Wang et al. 2001), which is then elaborated into separate but coordinated three-dimensional commands for eye and head rotation by the brainstem and cerebellum(Optican and Quaia 2002, Klier, Wang et al. 2003). Given that our analysis separates T and G based on endogenous variable gaze errors, this suggests that the SC (or a circuit that includes the SC) is involved in producing those errors. Conversely, we cannot conclude that our transformation result generalizes to all situations with different tasks and error types.

### What produces the TG transformation?

In this study we can only comment directly on SC data, but the sensorimotor transformations for gaze likely involve its reciprocal connections to the frontal eye fields, cerebellum, and thalamus, as well as feedback from the brainstem (Munoz and Guitton 1985, Schall and Thompson 1999, Optican and Quaia 2002, Schall 2002, Sommer and Wurtz 2002). It has been suggested that studies which separated sensory and motor produced a transformation by activating separate circuits of cells to code different spatial variables (Pierrot-Deseilligny, Rivaud et al. 1991, Gaymard, Ploner et al. 1999, Ohbayashi, Ohki et al. 2003, Bays, Gorgoraptis et al. 2011, Barber, Caffo et al. 2013, Sajad, Sadeh et al. 2016, Sajad A 2016). To test if this was also the case here, we compared overall spatial coding in visual (V), visumotor (VM), and motor (M) neurons, but concluded this had little direct influence on the spatial code in this particular task. This need not always the case: in the FEF we found that visuomotor and motor cells code different spatial attributes at the end of a memory delay (Sajad et al. 2016). At this time, it cannot be said whether this difference is due to the difference in brain structures, or different tasks. Further, based on our data we cannot exclude the possibility that some other cell classification scheme might explain spatial coding better, or that V, VM, and M cells might make different contributions to some other gaze task.

When viewed as a spatiotemporal transformation (Figures 7 and 8), it became clear that the main determining factor for the SC spatial code during the reactive task was timing. This was distributed throughout different cell types and was perhaps most surprising in cells that fell within our visual classification. The most likely explanation for this is that the SC is involved in a noisy, distributed sensorimotor transformation(Burns and Blohm 2010, Franklin and Wolpert 2011) that includes lateral and recurrent connections(Harting 1977, Harting, Huerta et al. 1980, Meredith and Stein 1983, Fries 1984, May 2006). In this scenario, a major component of variable gaze errors results from the rapid accumulation and general spread of noise during the transformation from visual inputs to motor outputs, and we see this reflected in all of our SC cells. This noise is relative small during normal gaze shifts, but could become quite large during certain clinical conditions (Ketcham, Hodgson et al. 2003, Rottschy, Kleiman et al. 2013, Avery and Krichmar 2015) For this reason, the analysis tools used here could be useful for detecting biomarkers of the source of sensorimotor function in the affected circuits.

### Conclusions

To our knowledge, this is the first study to track the spatiotemporal code in superior colliculus cells during simple reactive saccades toward a briefly flashed target, demonstrate a rapid visuomotor transformation, and trace this to the accumulation of errors in a distributed SC circuit rather than a relay between cells with fixed spatial codes. We cannot say if these results generalize to other brain areas, tasks, and motor behaviors, but given the relative simplicity of our task and the evolutionary conservation of SC function, it seems likely that similar processes occur alone or in conjunction with other transformations in many other areas and behaviors.

## Acknowledgements

This project was supported by a grant from the Canadian Institute of Health Research. J.D Crawford is supported by a Canada Research Chair award. M.Sadeh and A. Sajad were supported by Ontario Graduate Scholarships. The authors thank S.Sun for computer programming support.

## References

Avery, M. C. and J. L. Krichmar (2015) “Improper activation of D1 and D2 receptors leads to excess noise in prefrontal cortex.” Front Comput Neurosci 9: 31.

Basso, M. A. and R. H. Wurtz (1998) “Modulation of neuronal activity in superior colliculus by changes in target probability.” J Neurosci 18(18): 7519–7534.

Brown, M. R., J. F. DeSouza, H. C. Goltz, K. Ford, R. S. Menon, M. A. Goodale and S. Everling (2004) “Comparison of memory- and visually guided saccades using event-related fMRI.” J Neurophysiol 91(2): 873–889.

Burns, J. K. and G. Blohm (2010) “Multi-sensory weights depend on contextual noise in reference frame transformations.” Frontiers in human neuroscience 4: 221.

Crawford, J. D. and D. Guitton (1997) “Visual-motor transformations required for accurate and kinematically correct saccades.” J Neurophysiol 78(3): 1447–1467.

Crawford, J. D., D. Y. Henriques and W. P. Medendorp (2011) “Three-dimensional transformations for goal-directed action.” Annu Rev Neurosci 34: 309–331.

Everling, S., M. C. Dorris, R. M. Klein and D. P. Munoz (1999) “Role of primate superior colliculus in preparation and execution of anti-saccades and pro-saccades.” J Neurosci 19(7): 2740–2754.

Franklin, D. W. and D. M. Wolpert (2011) “Computational mechanisms of sensorimotor control.” Neuron 72(3): 425–442.

Freedman, E. G. and D. L. Sparks (1997) “Activity of cells in the deeper layers of the superior colliculus of the rhesus monkey: evidence for a gaze displacement command.” J Neurophysiol 78(3): 1669–1690.

Fries, W. (1984). “Cortical projections to the superior colliculus in the macaque monkey: a retrograde study using horseradish peroxidase.” J Comp Neurol 230(1): 55–76.

Gnadt, J. W., R. M. Bracewell and R. A. Andersen (1991) “Sensorimotor transformation during eye movements to remembered visual targets.” Vision Res 31(4): 693–715.

Goldman-Rakic, P. S. (1995). “Cellular basis of working memory.” Neuron 14(3): 477–485.

Harting, J. K. (1977). “Descending pathways from the superior colliculus: an autoradiographic analysis in the rhesus monkey (Macaca mulatta).” Journal of comparative neurology 173(3): 583–612.

Harting, J. K., M. F. Huerta, A. J. Frankfurter, N. L. Strominger and G. J. Royce (1980) “Ascending pathways from the monkey superior colliculus: an autoradiographic analysis.” Journal of Comparative neurology 192(4): 853–882.

Hollingworth, A. (2015). “Visual working memory modulates within-object metrics of saccade landing position.” Ann N Y Acad Sci 1339: 11–19.

Horwitz, G. D. and W. T. Newsome (1999) “Separate signals for target selection and movement specification in the superior colliculus.” Science 284(5417): 1158–1161.

Ketcham, C. J., T. L. Hodgson, C. Kennard and G. E. Stelmach (2003) “Memory-motor transformations are impaired in Parkinson’s disease.” Exp Brain Res 149(1): 30–39.

Klier, E. M., H. Wang and J. D. Crawford (2001) “The superior colliculus encodes gaze commands in retinal coordinates.” Nat Neurosci 4(6): 627–632.

Klier, E. M., H. Wang and J. D. Crawford (2003) “Three-dimensional eye-head coordination is implemented downstream from the superior colliculus.” J Neurophysiol 89(5): 2839–2853.

May, P. J. (2006). “The mammalian superior colliculus: laminar structure and connections.” Progress in brain research 151: 321–378.

Mays, L. E. and D. L. Sparks (1980) “Dissociation of visual and saccade-related responses in superior colliculus neurons.” J Neurophysiol 43(1): 207–232.

McPeek, R. M. and E. L. Keller (2004) “Deficits in saccade target selection after inactivation of superior colliculus.” Nat Neurosci 7(7): 757–763.

Meredith, M. A. and B. E. Stein (1983) “Interactions among converging sensory inputs in the superior colliculus.” Science 221(4608): 389–391.

Miller, E. K., C. A. Erickson and R. Desimone (1996) “Neural mechanisms of visual working memory in prefrontal cortex of the macaque.” J Neurosci 16(16): 5154–5167.

Miyashita, N. and O. Hikosaka (1996) “Minimal synaptic delay in the saccadic output pathway of the superior colliculus studied in awake monkey.” Exp Brain Res 112(2): 187–196.

Moschovakis, A. K., A. B. Karabelas and S. M. Highstein (1988) “Structure-function relationships in the primate superior colliculus. II. Morphological identity of presaccadic neurons.” J Neurophysiol 60(1): 263–302.

Munoz, D. and D. Guitton (1985) “Tectospinal neurons in the cat have discharges coding gaze position error.” Brain research 341(1): 184–188.

Optican, L. M. and C. Quaia (2002) “Distributed model of collicular and cerebellar function during saccades.” Ann N Y Acad Sci 956: 164–177.

Pouget, A. and L. H. Snyder (2000) “Computational approaches to sensorimotor transformations.” Nat Neurosci 3 Suppl: 1192–1198.

Rottschy, C., A. Kleiman, I. Dogan, R. Langner, S. Mirzazade, M. Kronenbuerger, C. Werner, N. J. Shah, J. B. Schulz, S. B. Eickhoff and K. Reetz (2013) “Diminished activation of motor working-memory networks in Parkinson’s disease.” PLoS One 8(4): e61786.

Sadeh, M., A. Sajad, H. Wang, X. Yan and J. D. Crawford (2015) “Spatial transformations between superior colliculus visual and motor response fields during head-unrestrained gaze shifts.” Eur J Neurosci 42(11): 2934–2951.

Sajad, A., M. Sadeh, G. P. Keith, X. Yan, H. Wang and J. D. Crawford (2015) “Visual-Motor Transformations Within Frontal Eye Fields During Head-Unrestrained Gaze Shifts in the Monkey.” Cereb Cortex 25(10): 3932–3952.

Sajad, A., M. Sadeh, X. Yan, H. Wang and J. D. Crawford (2016) “Transition from Target to Gaze Coding in Primate Frontal Eye Field during Memory Delay and Memory-Motor Transformation.” eNeuro 3(2).

Sajad A. S. M., Yan X, Wang H and Crawford JD. (2016). Time course for the accumulation of errors in the superior colliculus during memory-guided gaze shift. Society for Neuroscience. San Diego, CA.

Schall, J. D. (2002). “The neural selection and control of saccades by the frontal eye field.” Philos Trans R Soc Lond B Biol Sci 357(1424): 1073–1082.

Schall, J. D. and K. G. Thompson (1999) “Neural selection and control of visually guided eye movements.” Annu Rev Neurosci 22: 241–259.

Snyder, L. H. (2000). “Coordinate transformations for eye and arm movements in the brain.” Curr Opin Neurobiol 10(6): 747–754.

Sommer, M. A. and R. H. Wurtz (2002) “A pathway in primate brain for internal monitoring of movements.” Science 296(5572): 1480–1482.

Sparks, D. L. (1989). “The neural encoding of the location of targets for saccadic eye movements.” J Exp Biol 146: 195–207.

Sparks, D. L. (2002). “The brainstem control of saccadic eye movements.” Nat Rev Neurosci 3(12): 952-964.

Sparks, D. L. and R. Hartwich-Young (1989). “The deep layers of the superior colliculus.” Rev Oculomot Res 3: 213–255.

Sparks, D. L. and J. D. Porter (1983) “Spatial localization of saccade targets. II. Activity of superior colliculus neurons preceding compensatory saccades.” J Neurophysiol 49(1): 64–74.

Stanford, T. R. and D. L. Sparks (1994) “Systematic errors for saccades to remembered targets: evidence for a dissociation between saccade metrics and activity in the superior colliculus.” Vision Res 34(1): 93-106.

Waitzman, D. M., T. P. Ma, L. M. Optican and R. H. Wurtz (1988) “Superior colliculus neurons provide the saccadic motor error signal.” Exp Brain Res 72(3): 649–652.

Walker, M. F., E. J. Fitzgibbon and M. E. Goldberg (1995) “Neurons in the monkey superior colliculus predict the visual result of impending saccadic eye movements.” J Neurophysiol 73(5): 1988–2003.

White, J. M., D. L. Sparks and T. R. Stanford (1994) “Saccades to remembered target locations: an analysis of systematic and variable errors.” Vision Res 34(1): 79–92.

Wurtz, R. H. and J. E. Albano (1980) “Visual-motor function of the primate superior colliculus.” Annu Rev Neurosci 3: 189–226.

Wurtz, R. H. and C. W. Mohler (1976) “Organization of monkey superior colliculus: enhanced visual response of superficial layer cells.” J Neurophysiol 39(4): 745–765.

